# Endothelial Cell Serum and Glucocorticoid Regulated Kinase 1 (SGK1) Mediates Vascular Stiffening

**DOI:** 10.1101/2023.10.19.563167

**Authors:** Liping Zhang, Zhe Sun, Yan Yang, Austin Mack, Mackenna Rodgers, Annayya Aroor, Guanghong Jia, James R. Sowers, Michael A. Hill

## Abstract

**Background:** Excess dietary salt increases vascular stiffness in humans, especially in salt-sensitive populations. While we recently suggested that the endothelial sodium channel (EnNaC) contributes to salt-sensitivity related endothelial cell (EC) and arterial stiffening, mechanistic understanding is incomplete. This study thus aimed to explore the role of EC-serum and glucocorticoid regulated kinase 1 (SGK1), as a regulator of sodium channels, in EC and arterial stiffening.

**Methods and Results:** A mouse model of salt sensitivity-associated vascular stiffening was produced by subcutaneous implantation of slow-release deoxycorticosterone acetate (DOCA) pellets, with salt (1% NaCl, 0.2% KCl) administered via drinking water. Preliminary data showed that global SGK1 deletion caused significantly decreased blood pressure, EnNaC activity and aortic endothelium stiffness as compared to control mice following DOCA-salt treatment. To probe EC signaling pathways, selective deletion of EC-SGK1 was performed by cross-breeding cadherin 5-Cre mice with sgk1^flox/flox^ mice. DOCA-salt treated control mice had significantly increased blood pressure, EC and aortic stiffness in vivo and ex vivo, which were attenuated by EC-SGK1 deficiency. To demonstrate relevance to humans, human aortic ECs were cultured in the absence or presence of aldosterone and high salt with or without the SGK1 inhibitor, EMD638683 (10uM or 25uM). Treatment with aldosterone and high salt increased intrinsic stiffness of ECs, which was prevented by SGK1 inhibition. Further, the SGK1 inhibitor prevented aldosterone and high salt induced actin polymerization, a key mechanism in cellular stiffening.

**Conclusion:** EC-SGK1 mediates salt-sensitivity related EC and aortic stiffening by mechanisms appearing to involve regulation actin polymerization.

**Graphical Abstract:** 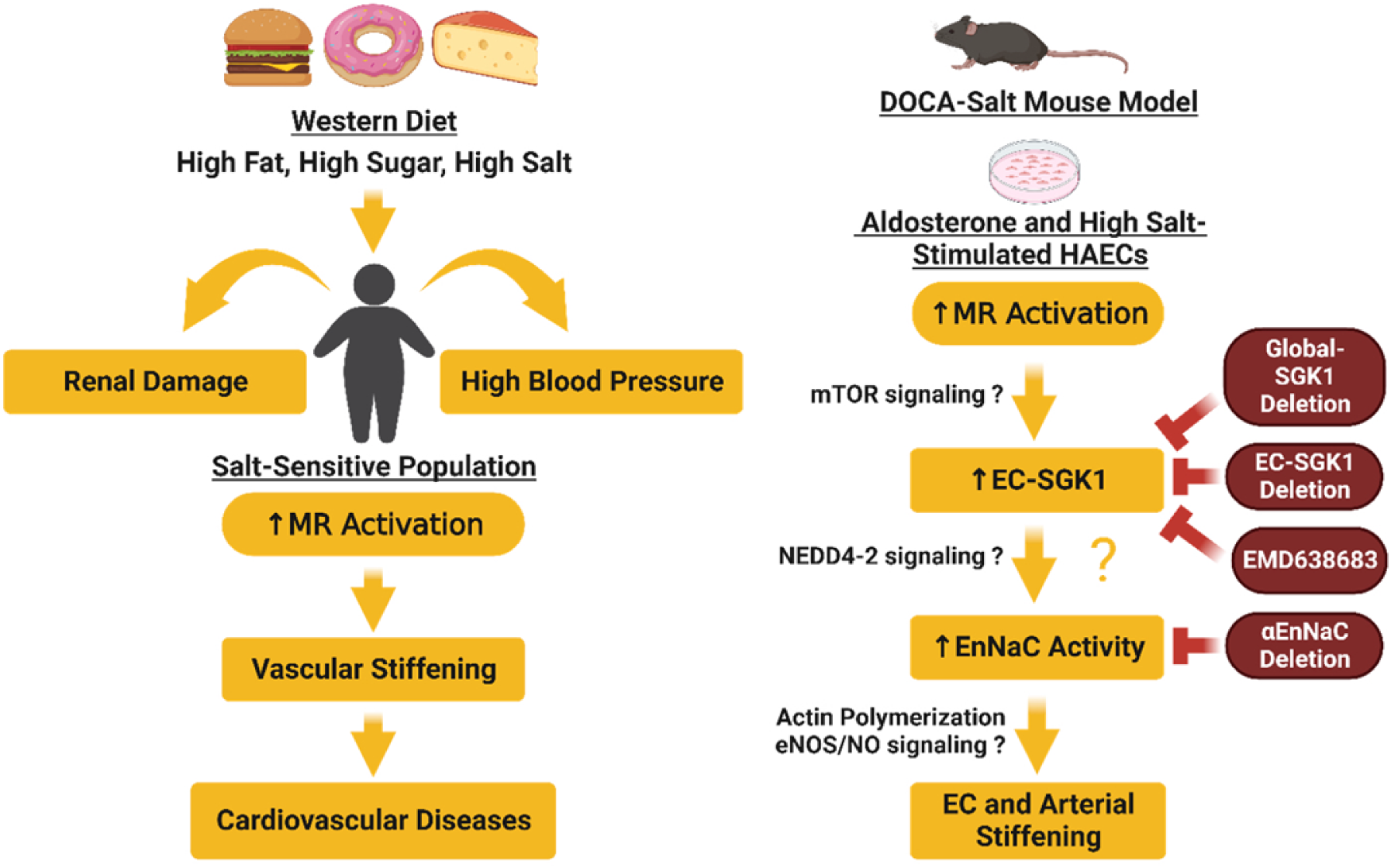

## Introduction

Cardiovascular disease remains the major cause of death, both in the United States ^1,2^ and globally ^3^. It is, thus, of great importance to detect cardiovascular disease in its early stages so that management with lifestyle changes and pharmacological interventions can be initiated before irreversible structural vascular changes occur. A considerable body of data exists to indicate that vascular stiffening, alone, is an independent predictor of future cardiovascular events in a variety of clinical situations including type 1 ^4^ and type 2 ^5^ diabetes, erectile dysfunction ^6^, coronary heart disease ^7^, chronic kidney disease ^8^, acute stroke ^9^, and resistant hypertension ^10^. A greater understanding of the mechanisms underlying vascular stiffening could, therefore, provide the rationale for novel approaches for diagnostic and therapeutic strategies for treatment of such cardiovascular-related pathologies.

A cross-sectional analysis of 723 children and young adults enrolled in a study to evaluate cardiovascular endpoints in obesity and type 2 diabetes mellitus has shown that sodium intake and sodium load are associated with increased arterial stiffness ^11^. Accumulating evidence in animal models of hypertension ^12,13^, as well as human patients with normal or elevated blood pressure ^14,15^, also support the association between dietary salt intake and arterial stiffening. Of note, high salt-induced increases in blood pressure and arterial stiffness were greater in individuals classified as being salt-sensitive ^14^. Interestingly, a recent study ^12^ which aimed to determine whether a high-salt diet further increases arterial blood pressure (ABP) and vascular dysfunction in a 2-kidney, 1-clip (2K1C) mouse model showed that systolic, diastolic, and mean ABP were significantly higher in mice fed 4.0% vs. 0.1% NaCl at 1 week but not after 3 weeks. These authors further suggested that this transient salt-sensitive hypertension might occur due to the peak activation of the renin-angiotensin-aldosterone-System (RAAS) at the early time-point.

A further study, using a cell culture approach, showed that a lack of aldosterone in the culture medium prevented high salt-induced increases in EC stiffness ^16^. These data together suggest both a mineralocorticoid, such as aldosterone, and salt are required to produce salt-sensitivity-related vascular dysfunction. Thus, in a recent study we utilized the deoxycortisone acetate (DOCA)-salt mouse model, which exhibits increased mineralocorticoid levels, replicating a situation of high salt consumption in salt-sensitive humans who do not manifest appropriate suppression of aldosterone production ^13^. The data showed that the combination of EC mineralocorticoid receptor (MR) activation and high salt consumption leads to activation of the endothelial sodium channel (EnNaC) together with increased EC and arterial stiffness and impaired EC-dependent vascular relaxation ^13^. Using a pharmacological intervention these studies further suggested that the underlying mechanisms might involve mTOR signaling ^13^. As it was previously shown in vivo that mTORC2 regulates renal tubule sodium uptake through serum and glucocorticoid regulated kinase 1 (SGK1)-dependent modulation of ENaC activity in C57BL/6 mice ^17^, we hypothesized that EC-SGK1 regulates salt-sensitivity-associated vascular stiffening through regulation of EnNaC.

SGK1, originally identified in rat mammary epithelial cells, is a transcriptional target of steroid hormones including glucocorticoids or mineralocorticoids (principally, aldosterone) in addition to other stimuli such as glucose and osmotic stress ^18^. Previous studies have suggested that SGK1 polymorphisms in humans (E8CC/CT; I6CC) associate with obesity ^19^ and diabetes ^20^. It has also been shown that constitutively active global SGK1 expression in mice exacerbates diet-induced obesity, metabolic syndrome, and hypertension ^18^. While few studies have examined the role of SGK1 in vascular stiffening, a recent study has shown that global SGK1 deficiency ameliorated smooth muscle cell calcification and stiffness during vitamin D3 overload–induced calcification ^21^. Accordingly, our preliminary data suggested global SGK1 deficiency attenuated DOCA-salt-induced aortic endothelium stiffening (Supplemental Figure S1). Nevertheless, a complete understanding of the role of SGK1 in ECs and salt-sensitivity-related vascular stiffening is lacking. Given that SGK1 is expressed in multiple tissues and cell types including smooth muscle cells (SMCs) ^22^, adipocytes ^23^, and macrophages ^24^, development of mouse models with EC-selective-SGK1 deficiency was proposed to aid in understanding the role of EC-SGK1 in vascular stiffening and potentially bring new mechanistic insights for developing more precise therapeutic interventions in salt-sensitivity-related vascular dysfunction.

## Materials and Methods

All animal procedures were performed in accordance with guidelines of the National Institutes of Health for the care and use of laboratory animals. All protocols were approved by the Animal Care and Use Committee of the University of Missouri-Columbia.

### Animals

5- to 7-month-old male and female mice with endothelial cell-selective deletion of the SGK1 (EC-SGK1^−/−^) and control littermates (EC-SGK1^+/+^) were generated by cross-breeding sgk1^flox/flox^ mice on a C57BL/6J genetic background with transgenic mice expressing Cre recombinase under the control of the Cadherin 5 promoter on a C57BL/6J and 129S1/SvImJ mixed genetic background (B6;129-Tg (Cdh5-cre) 1Spe/J). The DOCA-salt mouse model was generated by subcutaneous implantation of slow-release DOCA pellets (100 mg, Innovative Research of America, Sarasota, FL) and the addition of NaCl in the drinking water (1% sodium chloride with 0.2% potassium chloride) for 42 days. The dose and duration of this treatment was based on our previous observation that 50 mg/21 day-treatment of DOCA-salt only produced a mild phenotype of arterial stiffening and fibrosis ^13^. Potassium chloride was added to the drinking water to prevent DOCA-salt-induced decreases in serum potassium as reported in previous studies ^25,26^. The overall study timeline was shown in Supplemental Figure S2A. Three experimental groups were utilized as shown in Supplemental Figure S2B: EC-SGK1 knockout mice implanted with DOCA drinking salt water (DOCA-Salt EC-SGK1^-/-^); littermate control mice implanted with DOCA drinking salt water (DOCA-Salt EC-SGK1^+/+^); and littermate control mice underwent a sham implantation drinking tap water (Sham EC-SGK1^+/+^).

### Tail-cuff Blood Pressure Measurement

Blood pressure was measured using a non-invasive tail-cuff system (CODA-HT2; Kent Scientific, Torrington, CT) immediately prior to initiating treatments and at the end of the study. Mice were acclimatized to physical restraint and the tail-cuff measurement procedures for four consecutive days prior to blood pressure determination. All blood pressure measurements were conducted at the same time of the day (1 to 3PM) and by the same person to limit the influence of the circadian rhythm and reduce handling variability. A minimum of ten blood pressure readings were averaged for each animal to obtain final readings of systolic and diastolic pressures.

### Ultrasound Measurements

As an index of vascular stiffness, pulse wave velocity (PWV) was measured in the abdominal aorta using high frequency ultrasound (Vevo 2100, VisualSonics, Toronto, ON, Canada). In vivo ultrasound measurements were performed immediately prior to initiating treatments and at the end of the study using a previously published protocol ^27^. PWV is considered the gold standard to determine large artery stiffness, in vivo ^28^. As additional indicators of stiffness, aortic distensibility and aortic radial strain were subsequently calculated using Vevo LAB and Vevo Vasc software packages (Visualsonics, Toronto, ON, Canada). To enhance rigor, ultrasound measurements were performed with the operator blinded to the mouse strain or treatments.

### Atomic Force Microscopy

Aortic EC stiffness was measured in en face aortic segments using indentation atomic force microscopy (AFM) according to a previously published protocol ^29^. In brief, the thoracic aorta was freshly isolated, and the adjacent tissues were carefully removed under a dissection microscope. A 2 mm aortic ring was cut open longitudinally to allow access to the endothelial surface. The aortic explants were then glued to a coverslip using cell and tissue adhesive and transferred to a 60mm culture dish. The aortic explant was cultured in incubator at 37°C, 5% CO_2_ for 30 min before subsequent *en face* measurement of stiffness. Repeated cycles of nano- indentation and retraction cycles on the cell surface were interrogated using an AFM cantilever (MLCT, Bruker-nano, Goleta, CA).

### Primary Cell Isolation and PCR

Primary ECs, SMCs, and monocytes were isolated from EC-SGK1 knockout mice and their littermate control mice. For primary ECs isolation, fresh mouse lung tissues were collected and dissociated using a lung dissociation kit (Miltenyi Biotec Inc., Auburn, CA, USA). Lung ECs were then isolated using two step magnetic bead separation (CD45 and CD31 antibody-conjugated microbeads; Miltenyi Biotec Inc., Auburn, CA, USA). For primary SMCs isolation, mouse aorta was freshly collected into cold PSS buffer (145 mM NaCl, 4.7 mM KCl, 2 mM CaCl2, 1 mM MgSO4, 1.2 mM NaH2PO4, 0.02 mM EDTA). Adjacent tissues, the adventitia, and the endothelial layer of the aorta were carefully removed in a pre-chilled dissection chamber (4 °C). The aorta was then transferred to first digestion buffer (1 mg/ml DTE (dithioerythriotol), 27 unit/ml Papain in PSS), and incubated for 20 minutes at 37°C, followed by incubation in a second digestion buffer (0.975 unit/ml Collagenase H, 0.9 mg/ml Collagenase F, 1 mg/ml Soybean trypsin inhibitor in PSS) for 4 minutes. After incubation, the digestion buffer was carefully removed without disrupting the aorta and replaced with small amount of PSS buffer. The aorta was disrupted by pipetting (2 to 5 times) and centrifuged for 5 min at 300 g, 4 °C. The supernatant was discarded and SMCs were collected. For monocyte isolation, mouse spleens were freshly placed into PBS with 2% FBS and washed 2-3 times with PBS. Scissors were used to reduce the spleen into small pieces in a 60mm culture dish containing 1mL of PBS. Subsequently, the homogenate was transferred to a 70um filter positioned over a 60mm culture dish to collect the sample. Spleen homogenates were then further disrupted by trituration using a 3mL syringe plunger. The homogenate was then transferred to a 30um filter positioned above a 15mL centrifuge tube to collect the single cell suspension sample. Monocytes were then isolated using the EasySep™ mouse monocyte isolation kit following the instruction manual. Cell samples were then prepared for rtPCR for measurement of sgk1 mRNA expression.

### Cell Culture

Primary human aortic endothelial cells (HAECs, Cat. No. 6100, ScienCell Research Laboratory, Carlsbad, CA) were grown in endothelial cell culture medium (ECM, Cat. No. 1001, ScienCell) with endothelial cell growth supplement (ECGS, Cat. No. 1052, ScienCell), antibiotic solution (P/S, Cat. No. 0503, ScienCell), and 5% charcoal stripped FBS (Cat. No. A3382101, Gibco, Detroit, MI) at 37°C, 5% CO2 incubator. All cell culture vessels including T75 flask, 60mm dishes, 6 well plate, and Ibidi 15 well chambered slides (Ibidi, Gräfelfing, Germany) were coated with gelatin-based coating solution (Cat. No. 6950, CellBiologics, Chicago, IL) before seeding cells. HAECs were used at passages 2 to 5. Once reaching 90% confluence, HAECs were exposed to the experimental treatments. To induce cell stiffening, HAECs were treated with aldosterone (10^-7^ M) and added NaCl (+16 mM) for 48 hours. To pharmacologically inhibit SGK1 activity, additional cells were pretreated with either 10 μM or 25 μM of the selective SGK inhibitor, EMD638683 (Cat. No. A3389, APExBIO, Houston, TX) for 1h before being subjected to 48 h treatment of aldosterone and salt. To exclude the potential effect of changes in osmolarity, additional HAECs were treated with mannitol (32 mM) and served as the control. Further, the amount of DMSO, used as a diluent for some chemicals, was adjusted to be the same in all treatment groups.

### Immunofluorescence Staining and Image Analysis

HAECs were seeded on a 15 well chambered slides (Ibidi, Gräfelfing, Germany) and incubated with respective treatments as mentioned previously. After 48 h treatment, cells were rinsed twice with PBS and fixed with 2% paraformaldehyde for 15 min at room temperature. Fixed cells were washed once with 0.1 M glycine to quench remaining paraformaldehyde and twice with PBS for 5 min. Cells were then permeabilized with 0.1% Triton X-100 for 10 min following by 2×5 min washes with PBS. Next, cells were stained with Alexa-568 Phalloidin (1:400) and Alexa-488 Deoxyribonuclease I (1:200, 25 μg/mL) for 1 h at room temperature while protected from light. After staining, cells were washed with PBS (3×5min) and mounted using ProLong™ Glass Antifade Mountant with NucBlue™ Stain. Images were acquired on a confocal microscope (Leica TCS SPE, Leica Microsystems Inc., Deerfield, IL) using a 20X/0.60 objective. Since immunofluorescence staining is considered semi-quantitative, several steps were taken to minimize variation in staining, image acquisition and analysis. Staining protocols were kept the same for all the groups. To assure the specificity of our measurement for actin polymerization, additional cells were treated with Cytochalasin D (10 μM), a reagent that disrupts actin polymerization and serves as a negative control. The settings of the imaging system, such as laser power, gain, and zoom factors, were kept constant across all groups. Fluorescence signals were carefully monitored to avoid image saturation. In addition, to avoid bias in image analysis and to eliminate the effect of cell density on fluorescence signal, we used a tailor-made MatLab script in which whole images were divided by regions of interest (ROI) of 200×200 pixels (25 ROIs in total), and the mean intensities of these ROIs for each channel were calculated. The ten ROIs with the highest mean intensity for F-actin were mathematically identified in a blinded fashion, and the average of these ten ROIs was used as a single value for statistical analysis. G-actin signals within these ROIs were also determined and averaged.

### Statistical Analysis

Data are presented as mean ± SEM. Differences between two treatment groups were determined using unpaired Student t-tests; differences between three or four treatment groups were determined using one-way ANOVA with Tukey’s multiple comparisons analysis or two-way ANOVA with Sidak’s multiple comparisons after passing normality testing. When data did not conform to a Gaussian distribution, raw data Y was transformed to Log (Y) before comparison analysis. A value of P<0.05 was considered statistically significant. Figures were generated and statistical analyses performed using GraphPad Prism 8.0 software.

## Results

### Validation of EC-SGK1 knockout mouse model

The EC-SGK1 mouse model was successfully produced by crossing the Cre mouse carried an EC-specific promoter (Cadherin 5) with sgk1^flox/flox^ mouse (Figure 1A). Representative genotyping gels showed that EC-SGK1 knockout mice expressed both the Cadherin 5-Cre and sgk1-floxed allele while the EC-SGK1 control mice only expressed sgk1-floxed allele but not Cre. The negative control wildtype mice showed no expression for both alleles (Supplemental Figure S3A). To validate that the deletion of SGK1 is selective for ECs, we isolated ECs, SMCs and monocytes from both the EC-SGK1 knockout and littermate control mice. The data showed that SGK1 mRNA levels were significantly reduced in ECs, but importantly not in vascular SMCs isolated from EC-SGK1 knockout mice as compared to those from littermate control mice (Figure 1B and Supplemental Figure S3B). mRNA expression of SGK1 was also downregulated in monocytes isolated from EC-SGK1 knockout mice, indicating non-specific activation of Cre in monocytes (Supplemental Figure S3C) consistent with the literature ^30,31^. Of note, there was no significant differences in blood pressure or aortic stiffness when comparing EC-SGK1^-/-^ mice with EC-SGK1^+/+^ mice at baseline level (Supplemental Figure S4).

**Figure 1.**
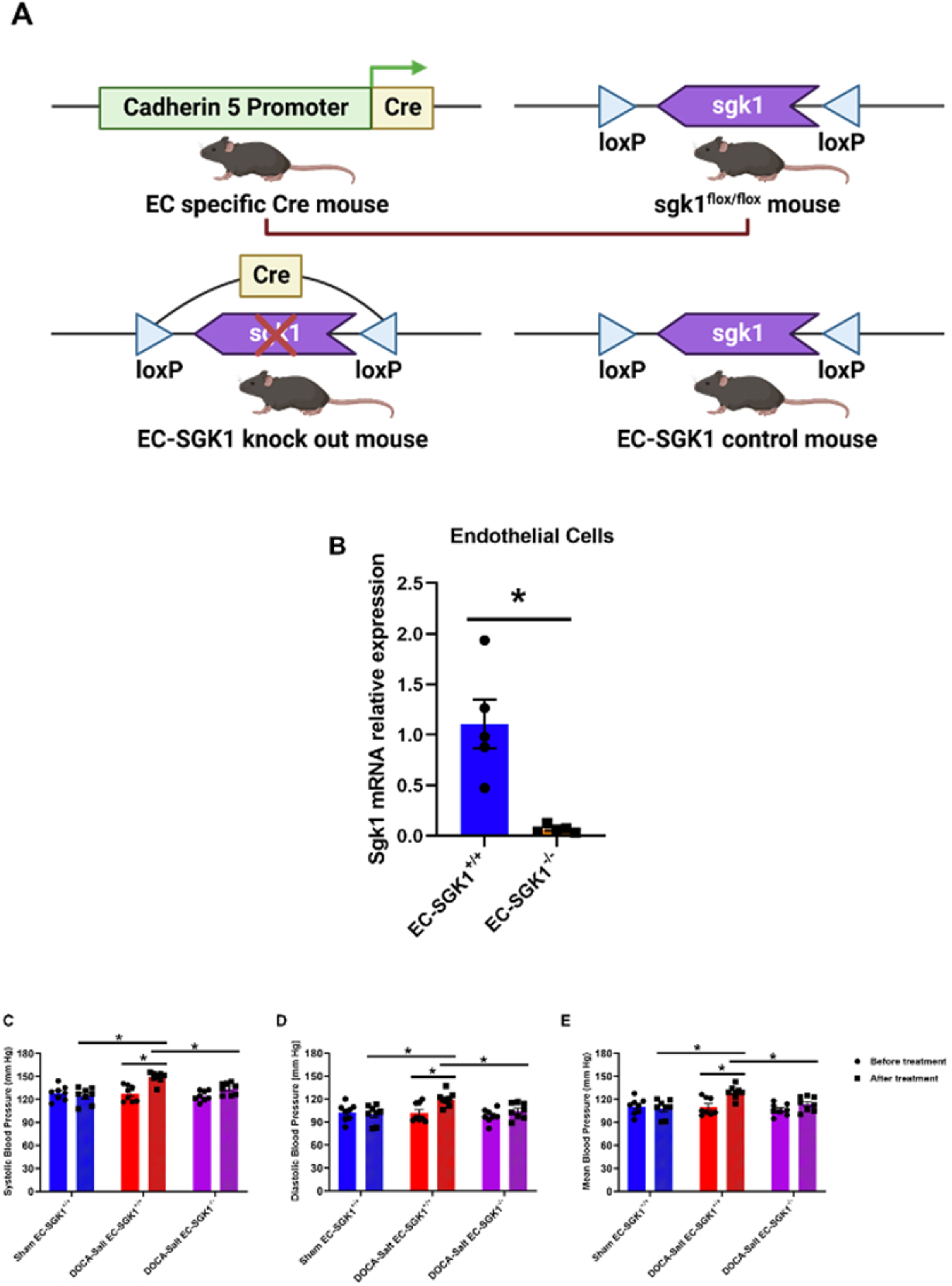
DOCA-salt-induced increases in blood pressure were prevented by EC-SGK1 deficiency. (A) Schematic diagram of the breeding protocol. (B) Sgk1 mRNA relative expression in endothelial cells. (C) Systolic blood pressure, (D) Diastolic blood pressure, and (E) Mean blood pressure measured before and at the completion of each treatment n=8 mice per treatment group. * P<0.05, using two-way ANOVA with Sidak’s multiple comparison analysis. N.S. indicates no significant differences.

### Characteristics of animal treatment groups

As there were no obvious sex differences in BP, ultrasound or AFM measurements, female and male data have been combined. Consistent with previous observations ^32,33^, plasma sodium levels were significantly increased while plasma potassium levels were significantly decreased in all DOCA-salt-treated groups (Table S1). The decreases in plasma potassium suggests that 0.2% potassium chloride supplement in drinking water was not sufficient to totally compensate the potassium loss due to the 42-day-DOCA-salt administration, nevertheless, potassium levels in all groups remained within the physiological range (3.5 to 5 mEq/L) ^34^. In addition, there were no significant differences in plasma glucose, insulin, or homeostatic model assessment-insulin resistance (HOMA-IR) among the groups (Supplemental Table S1). Of note, all DOCA-salt treated groups had slightly increased body weight during the first week post treatment and exhibited polydipsia during the whole treatment period (Supplemental Figure S5A-B). On the day of tissue harvest, fasting (6 hours) bodyweight normalized to tibia length showed no significant differences among all of the treatment groups (Supplemental Figure S5C). Isolated heart, left and right kidney, as well as liver weights were significantly increased while epididymal fat were significantly decreased in DOCA-salt-treated animals (Supplemental Figure S5D-E), suggesting that 42 day-treatments of DOCA-salt might affect overall body fat distribution and induces hypertrophy in heart, kidney and liver. It is well documented that DOCA-Salt administration induces both cardiac ^26,35,36^ and renal hypertrophy and fibrosis ^26,37^. Indeed, a previous study ^38^ has shown that 12 weeks of DOCA-salt administration induced marked glomerular sclerosis and tubulointerstitial damage with interstitial fibrosis and inflammation in kidneys, which were prevented in mice with global SGK1 deletion. Interestingly, our data showed that EC-SGK1 deletion attenuated renal hypertrophy, suggesting a potential role for EC-SGK1 in the renal pathology associated with DOCA-salt administration.

### DOCA-salt-induced increases in blood pressure were prevented by EC-SGK1 deficiency

After the 42-day treatment period, DOCA-salt administration significantly increased systolic blood pressure (SBP), diastolic blood pressure (DBP), and mean blood pressure (MBP) in EC-SGK1^+/+^ mice, which were prevented by EC-SGK1 deficiency (Figure 1 C-E).

### DOCA-salt-induced increases in arterial and aortic intimal cellular stiffness were attenuated by EC-SGK1 deficiency

To evaluate arterial stiffness, high frequency ultrasound measurements were performed on all experimental groups immediately prior to and at the end of the respective treatments. Our data showed an increased aortic PWV in both the EC-SGK1^+/+^ mice and EC-SGK1^-/-^ mice when comparing pre to post treatment values in individual treatment group, indicating DOCA-salt increased arterial stiffness. Importantly, there was a significant decrease in PWV when comparing EC-SGK1^-/-^ mice and EC-SGK1^+/+^ mice post treatment values, indicating that DOCA-salt-induced arterial stiffening was indeed attenuated by EC-SGK1 deficiency (Figure 2A). We further evaluated stiffness of the endothelium by en face indentation AFM using freshly prepared aortic explants. The data showed that endothelial stiffness was increased in DOCA-salt treated EC-SGK1^+/+^ mice when compared to sham EC-SGK1^+/+^ mice. Importantly, the DOCA-salt induced increase in aortic endothelium stiffness was prevented by EC-SGK1 deficiency (Figure 2B). Together, these data suggest that EC-SGK1 mediates DOCA-salt-induced vascular stiffening in mice.

**Figure 2.**
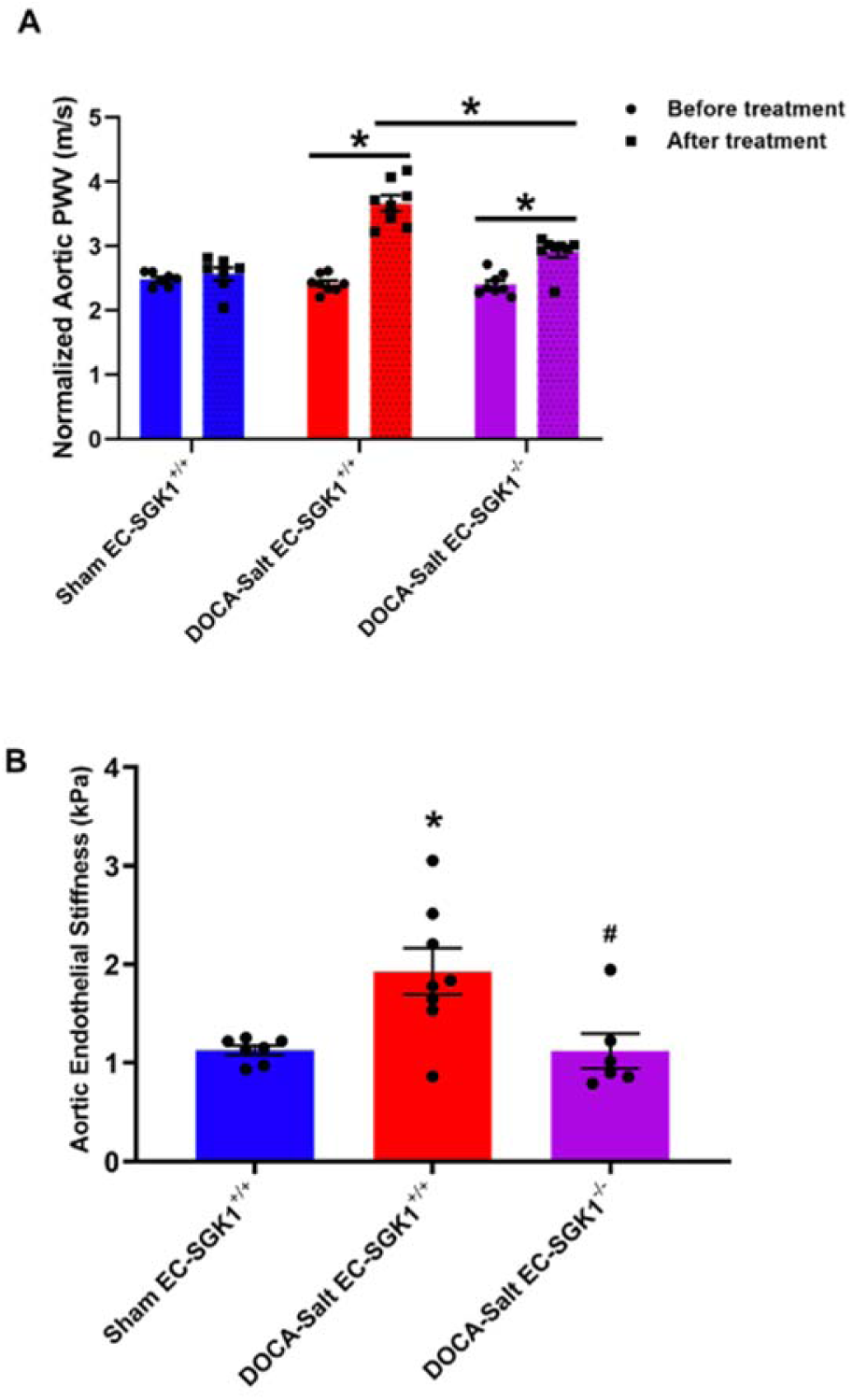
EC-SGK1 deficiency attenuated DOCA-salt-induced increase in arterial and aortic endothelium stiffness. (A) Aortic pulse wave velocity measured in vivo. (B) Ex vivo, in situ, aortic endothelium stiffness in arterial explants measured by atomic force microscopy. n=6- 8 mice per treatment group. For ultrasound data, * P<0.05, using two-way ANOVA with Sidak’s multiple comparison analysis. For atomic force microscopy data, * P<0.05, compared to Sham EC-SGK1^+/+^; # p<0.05, compared to DOCA-Salt EC-SGK1^+/+^ using one-way ANOVA with Tukey’s multiple comparison analysis.

### Aldosterone and high salt induced HAECs stiffening was prevented by SGK1 inhibition

To determine if our findings in animal studies could be replicated in human cells, we initiated studies to test the role of EC-SGK1 in primary human aortic endothelial cells (HAECs). Consistent with previous literature ^16^, our data showed that exposure (48hrs) to both aldosterone (10^-7^M) and salt (+ 16 mM NaCl) induced EC stiffening (Figure 3). Importantly, both a relatively low dose (10 μM) and a high dose (25 μM) of a selective SGK1 pharmacological inhibitor (EMD638683) prevented the increased stiffness in HAECs (Figure 3A), suggesting a role of SGK1 in regulation of EC stiffening in a model relevant to human disease.

**Figure 3.**
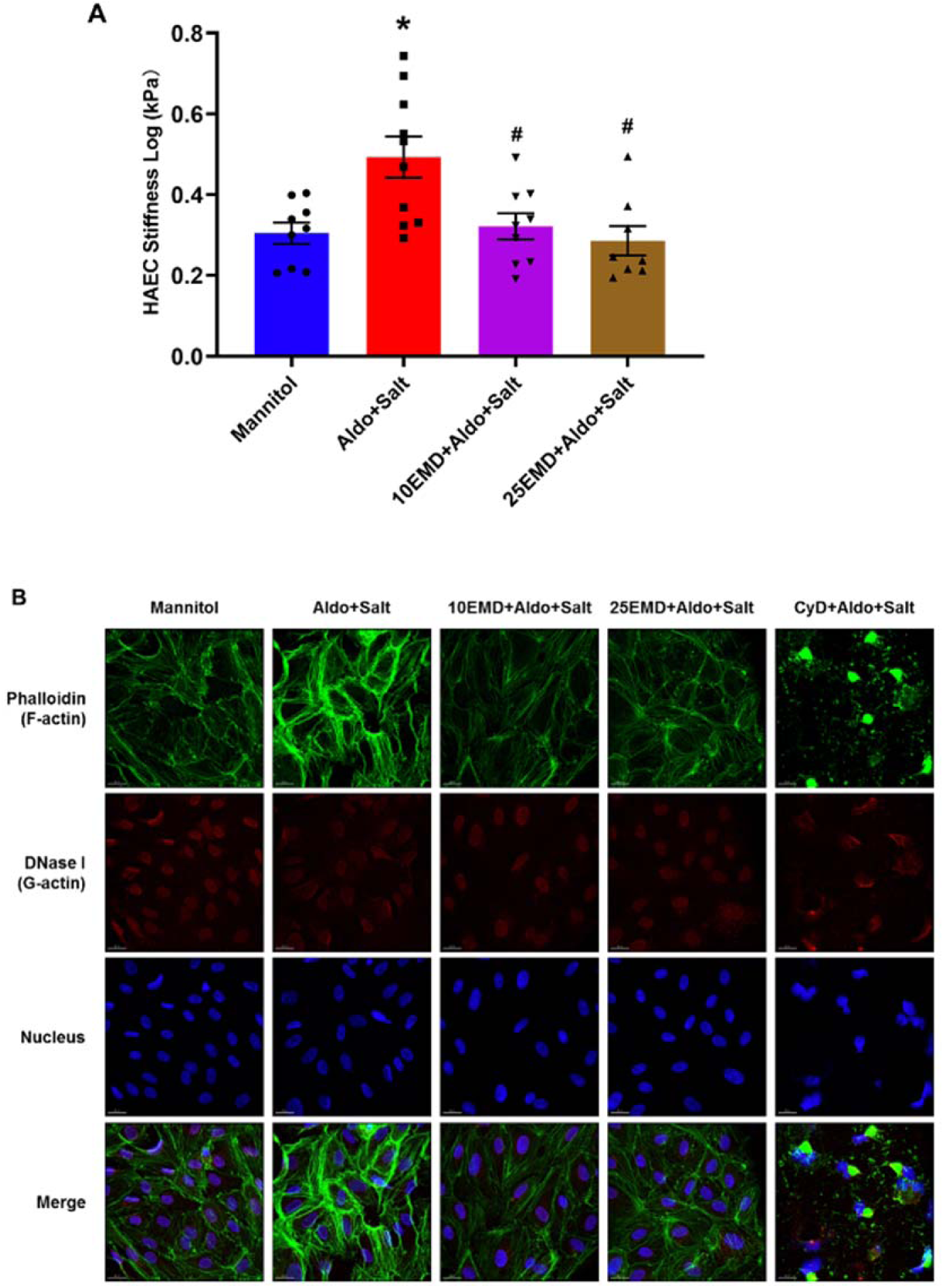
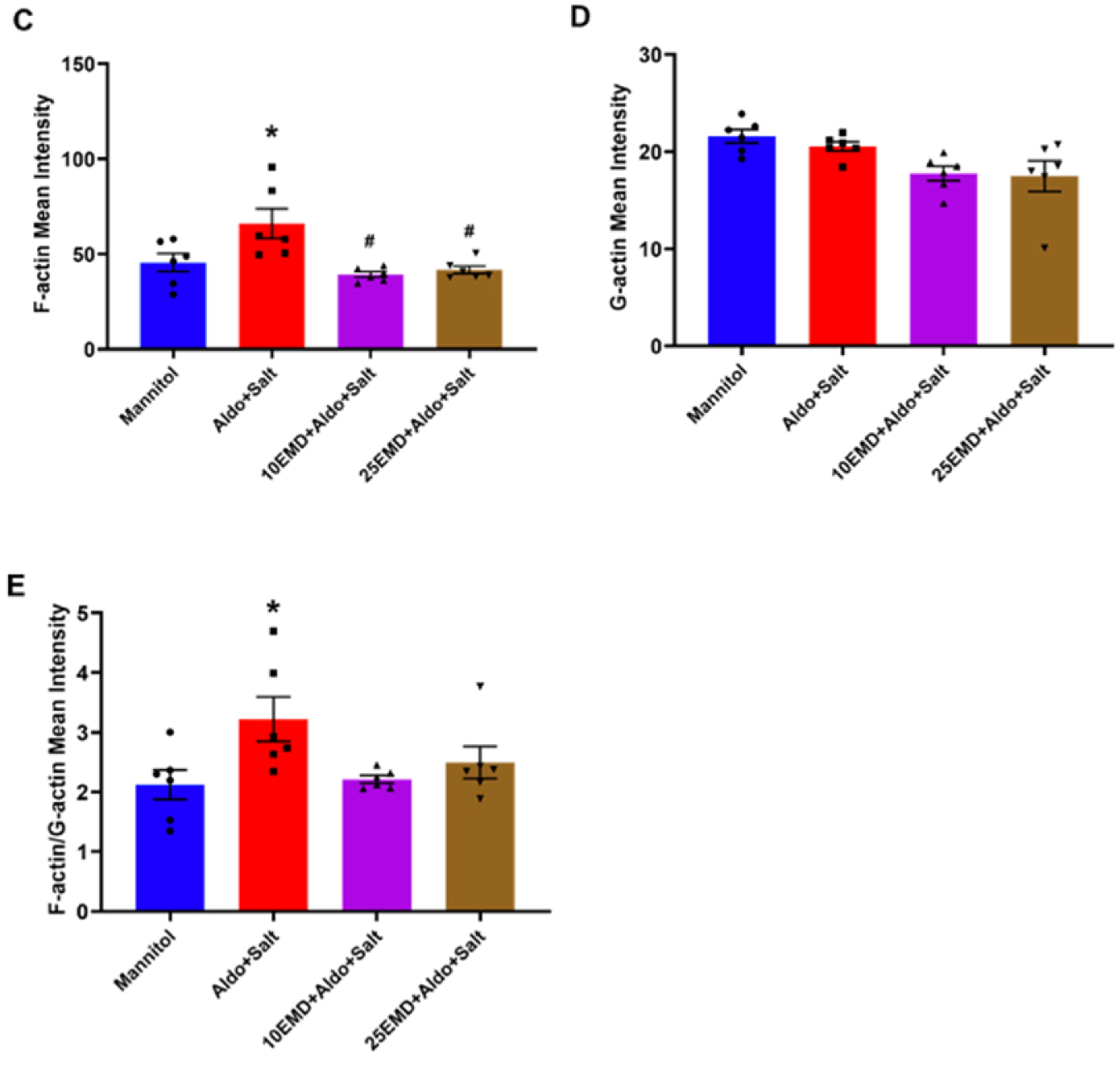
Aldosterone and high salt-induced stiffening of HAECs were prevented by SGK1 inhibition. (A) Endothelium stiffness measured by atomic force microscopy. (B) Representative images of F-actin, G-actin, and Nucleus staining on HAECs. (C) Quantification of F-actin and (D) G-actin mean intensity as well as (E) F-actin/G-actin ratio. n=8-10 per treatment group for atomic force microscopy; n=6 per treatment group for immunofluorescence staining. * P<0.05, compared to Mannitol; # p<0.05, compared to Aldo+Salt using one-way ANOVA with Tukey’s multiple comparison analysis.

### Aldosterone and high salt induced actin polymerization was prevented by SGK1 inhibition

To identify contributing cellular and molecular mechanisms underlying HAEC stiffening, we examined the effect of aldosterone and salt on actin polymerization in HAECs by comparing F-actin and G-actin expression. Our data showed that F-actin was significantly increased in aldosterone and salt treated groups, which was prevented in both groups treated by pharmacological SGK1 inhibition (Figure 3B). In contrast, there was no significant differences in G-actin expression among the treatment groups (Figure 3C). F-actin to G-actin ratio was significantly increased in the aldosterone and salt treated group but not SGK1 inhibitor treated groups as compared to the mannitol control group (Figure 3E). In addition, as a negative control, Cytochalasin D disrupted actin polymerization and showed aggregates/clusters formation around the nucleus (Figure 3A).

## Discussion

The epithelial sodium channel (ENaC) and SGK1 are both associated with blood pressure regulation. Indeed, Liddle’s syndrome ^39^, a form of hypertension, is caused by an ENaC gain of function gene variant ^40^. Similarly, people with polymorphisms in SGK-1 have been shown to have increased risk of hypertension and were more sensitive to the blood pressure elevating effects associated with hyperinsulinemia ^41^. Earlier studies on ENaC, SGK1, and their relationship have thus been largely focused on renal function and blood pressure regulation. In the past two decades, the role of the endothelial cell form of ENaC (EnNaC) in vascular dysfunction, particularly in females with obesity has been gradually identified ^29,42,43^, based on studies of Oberleithner et al. who initiated a focus on the role of EnNaC in the vascular endothelium ^44–47^. Despite this, direct regulators, and downstream targets of EnNaC that mediate vascular dysfunction, especially in relation to salt-sensitivity, are still not fully understood.

In the current study, we investigated the role of SGK1, a potential upstream regulator of EnNaC, in salt-sensitivity related vascular stiffening. In preliminary studies we found that while DOCA-salt treatment induced a significant increase of SBP in littermate control mice, SBP in mice with global SGK1 deletion remained unchanged after DOCA-salt treatment (Supplemental Figure S1A). Further, global SGK1 deletion attenuated DOCA-salt induced increases in aortic endothelium stiffening measured by AFM (Supplemental Figure S1D). However, aortic PWV measurement showed no significant differences in both the littermate control and global SGK1 knockout mice when comparing before versus after treatment (Supplemental Figure S1E), possibly resulting from the comparatively short duration of this preliminary study (50 mg DOCA, 21 days) which may have lessened the development of marked aortic stiffness and that such subtle changes in stiffness might not be detectable in vivo. As a result of these observations, we chose to use a longer duration (100 mg DOCA, 42 days) for the current study involving the EC-SGK1^-/-^ mice. After 42 days of administration of DOCA-salt, littermate control animals exhibited increases in blood pressure, arterial stiffening as measured by PWV and aortic EC stiffening as measured by AFM, which were prevented/attenuated in EC-SGK1^-/-^ mice. Further, using human primary aortic ECs, we also found that treatment of a SGK1 inhibitor prevented aldosterone and salt-induced EC stiffening. Taken together, our findings from both animal studies using mice with EC-SGK1 deficiency and HAECs treated with a SGK1 inhibitor suggested that EC-SGK1 mediates salt-sensitivity related EC and vascular stiffening.

Importantly, our preliminary data also showed that EnNaC currents were significantly reduced in ECs isolated from DOCA-salt-treated global SGK1 knockout mice as compared to DOCA-salt-treated control animals (Supplemental Figure S1B), suggesting a direct link between SGK1 and EnNaC. EnNaC activity can be modulated in several ways including 1) number of channels on the plasma membrane through regulation of gene/protein synthesis, 2) open probability, 3) channel conductance, 4) trafficking to membrane, and 5) ubiquitination which led to internalization and degradation. Such regulatory mechanism may involve transcriptionally regulated events or/and acute post-translational modifications (e.g., phosphorylation). In this regard, the ubiquitin ligase neuronal cell expressed developmentally downregulated 4-2 (NEDD4-2) is associated with ubiquitination of ENaC ^48^. Importantly, SGK1 has been shown to regulate ENaC activity through NEDD4-2 in Xenopus oocytes ^19^, COS7 kidney cells ^49^, mouse collecting duct cell line ^50^, and the heart ^51^. Therefore, it is hypothesized that EC- SGK1mediated increases in EnNaC activity might occur through regulation of NEDD4-2 (Figure 4 and Supplemental Figure S6C). In this regard, our data (Figure S6B) showed that 48 h treatment of 10 uM or 25 uM EMD638683 (SGK1 inhibitor) is sufficient to inhibit SGK1 activity as shown by SGK1-dependent-phosphorylated and total form of N-Myc Downstream Regulated Gene family 1(NDRG1) ^17,52^. Importantly, SGK1 inhibitor pre-treated groups exhibited reduced phosphorylated Nedd4-2 when compared to aldosterone and salt treated group (Supplemental Figure S6C). Nevertheless, it appeared that there might be some variation in baseline activity between cell passages, and batch to batch of cell samples. Future studies thus require critical control of cell passage numbers, and to increase rigor by studying additional cell isolates (cells collected from different donors) to detect true differences.

**Figure 4.**
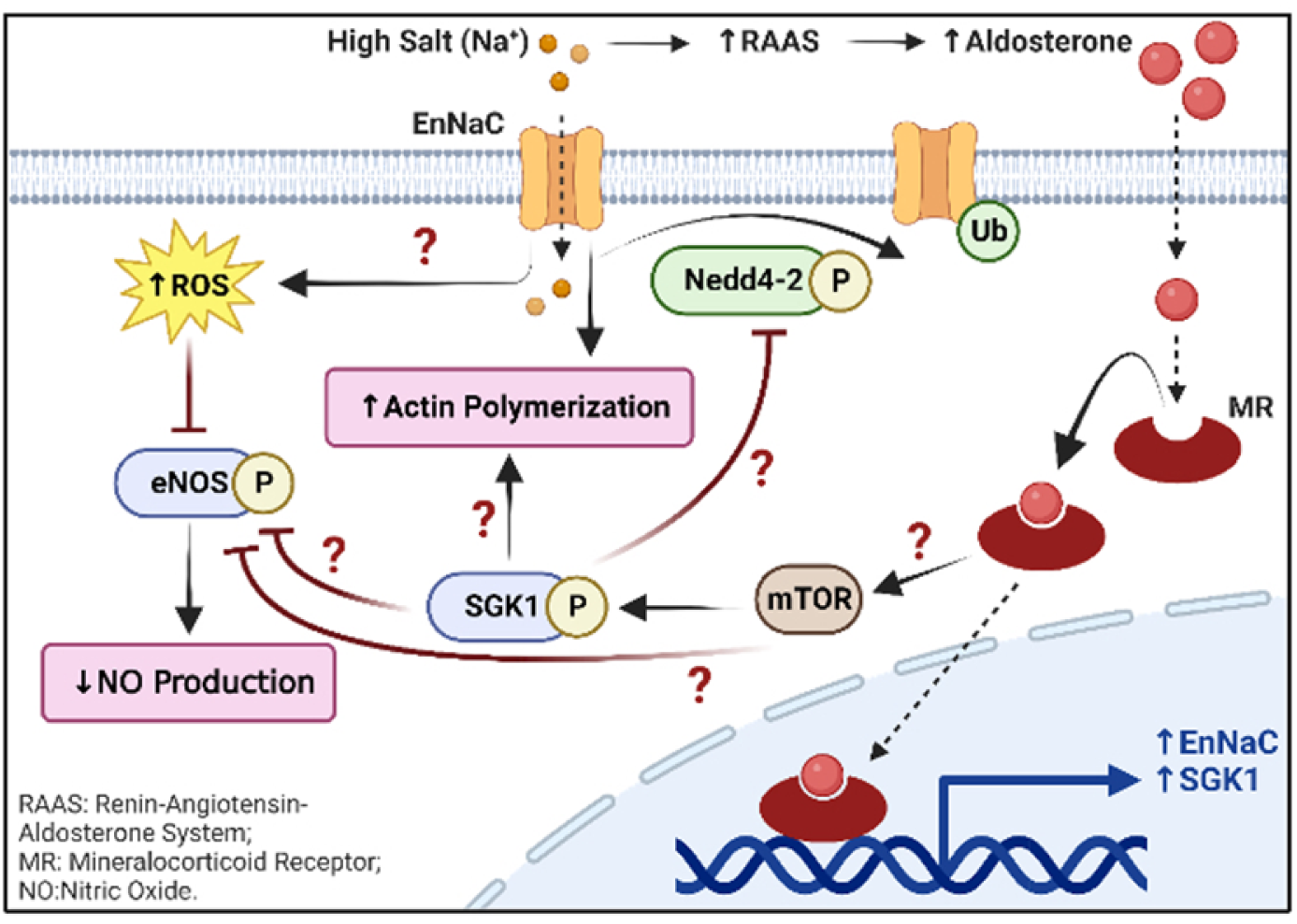
Proposed molecular signaling pathway. Aldosterone and high salt-induced stiffening of HAECs were prevented by SGK1 inhibition.

In terms of the mechanistic changes underlying EC stiffening in the current study, our data supported an important role for actin polymerization. Previous studies have shown that actin stress fiber formation stiffened the cell center of human umbilical vein endothelial cells ^53^. A further study demonstrated that high glucose significantly decreased release of NO along with stiffening of endothelial cell membranes and F-actin rearrangement ^54^, indicating a role of actin cytoskeleton in mediating EC stiffening. Importantly, the actin cytoskeleton has been shown to interact with various receptors, transporters, and ion channels ^55^. Indeed, Mazzochi et al. has shown that actin cytoskeleton directly binds to the carboxyl terminus of αENaC ^56^. Other studies suggest that ENaC may in turn affect cortical actin rearrangement by showing that rapid laminar shear stress led to a rapid MR-dependent membrane insertion of ENaC and subsequent stiffening of the endothelial cortex caused by actin polymerization ^57^. On the other hand, previous studies in mesenchymal stem cells showed that dexamethasone-induced cellular tension required SGK1 activation, which was involved in formation of fibronectin fibrils and their attachment to integrins at adhesion sites ^58^. In the current study, we proposed EC-SGK1 activation might lead to increased EnNaC activity and result in subsequent actin polymerization and EC cortical stiffening. Interestingly, a recent study^59^ related to Alzheimer’s disease (AD) suggested that hippocampal overexpression of SGK1 ameliorates cytoskeleton polymerization in middle-aged APP/PS1 mice while another earlier study ^60^ suggested that SGK1 regulates reorganization of actin cytoskeleton in mast cells upon degranulation. The controversy in the literatures suggested SGK1 might exert differential effects on actin dynamic in different cell types, further studies are thus required to examine the exact mechanisms underlying EC-SGK1 mediated actin polymerization.

In addition to actin dynamics, it is hypothesized that ECs might also mediate vascular stiffness through regulation of oxidative stress and endothelial nitric oxide (eNOS) production (Figure 4). Preventing endothelial dysfunction by improving oxidative stress and increasing endothelial NO production is regarded as a treatment strategy for a number of cardiovascular diseases ^61^. While studies have indicated that SGK1 can alter iNOS-mediated production of NO in renal epithelial cells ^62^, the role of EC-SGK1 in eNOS/NO signaling pathways and oxidative stress is yet to be examined in detail. In this regard, we observed significant increases in expression of total eNOS in the SGK1 inhibitor treated groups as compared to the aldosterone and salt treated groups, which suggests SGK1 regulation of eNOS expression (Figure S6D-E). Future studies using treatment with ROS scavengers or NOS inhibitors such as L-NAME might help us gain deeper insight in understanding the role of ROS/eNOS/NO signaling pathways in EC-SGK1 and EnNaC mediated vascular stiffness.

To our knowledge, this is the first study using mice with EC-SGK1 deficiency. This mouse model was produced using the Cre/LoxP system, a common genetic tool to control site specific recombination events by restricting the expression of Cre recombinase with a specific promoter, which then allows generation of experimental animals with tissue-specific genetic loss or gain of function. It should be noted, however, successful gene targeting with Cre/loxP critically depends on the precise expression pattern of the Cre recombinase. Payne et al. suggested that many EC-specific Cre have some degree of unreported activity, inconsistent recombination efficiency and/or parent-of-origin effects, which occur when the phenotypic effect of an allele depends on whether the allele is inherited maternally or paternally ^30^. The most common mouse model using a Cre promoter in ECs is the Tie2-Cre (also known as Tek-Cre). It is noted that Tie2-Cre mouse alleles are prone to variable and non-specific Cre recombinase activity and that all Tie2-Cre mouse models show some degree of Cre recombinase activity in the hematopoietic lineage ^30,63^. For example, by characterizing the Tie2-Cre;Rosa26R-EYFP reporter mice, Tang et al. ^64^ revealed Tie2-Cre-mediated recombination in 85% of the bone marrow cells population. Further, after analyzing the sub-classes of early hematopoietic progenitors, such as T cells, monocytes, granulocytes, and B cells, they found that ∼84% of each lineage was EYFP^+^, suggesting widespread contribution of Tie2-Cre to definitive hematopoietic cells. Similar to Tie2-Cre, the Cdh5-Cre alleles also showed Cre recombinase activity in hematopoietic cells. Indeed, Chen et al. ^31^ generated a VE-Cadherin-Cre transgene and used R26R-lacZ and R26R- YFP reporter mice to characterize the Cre activity. Their data has shown that 85% of fetal liver blood cells and 96% of CD45^+^ adult bone marrow cells were YFP^+^, suggesting unspecific Cre activity in the hematopoietic lineage. In the current study, we aimed to produce a mouse model with EC selective deletion of SGK1 not impacting other major cellular components of the blood vessel wall. Aware of potential unspecific Cre activity, we characterized our mice model by examining mRNA expression of SGK1 in isolated ECs, VSMCs (accounting for the major cellular components of the aorta), and monocytes in both Cdh5-Cre-positive;sgk1^flox/flox^ mice and Cadh5-Cre-negative;sgk1^flox/flox^ mice. Our data suggested nonspecific Cadh5-Cre activity in monocytes to be a limitation of the current study. Of note, to avoid potential side effect and toxicity of tamoxifen, the current study used a non-inducible Cre, future studies utilizing a tamoxifen-inducible Cre might or might not produce more precise expression of Cre in ECs. Further, using a Cre reporter gene such as GFP, YFP, and lacZ, or more reliable genetic tool such as the iSuRe-Cre ^65^ to monitor the spatial information on Cre recombinase activity is crucial given the unwanted Cre activity in tissue not of interests. Although we cannot exclude the potential effect of SGK1 deficiency in monocytes on our findings collected from the EC-SGK1 knockout mice, importantly, our in vitro studies using HAECs confirmed the important role of EC-SGK1 in EC stiffening, which is consistent with the animal experimental data.

The physiological mechanisms determining and regulating the onset and development of salt-sensitive hypertension are complex. Accumulating evidence has indicated genetic predisposition, biological sex and sex hormones, in combination with other mechanisms involving the RAAS, the endothelin system, nitric oxide (NO) and oxidative stress, the sympathetic nervous system, atrial natriuretic peptides (ANPs), CYP450-derived metabolites of arachidonic acid, and hyperinsulinemia ^66–68^. Human studies have suggested that polymorphisms of SGK1 and ENaC confer predisposition to hypertension, especially salt-sensitive hypertension ^41,69,70^. Consistent with these observations, our previous ^13^ and current study revealed a role of EnNaC and EC-SGK1 in salt-sensitive hypertension, respectively. Nevertheless, whether the effects of EnNaC and EC-SGK1 on BP regulation were through renal or circulatory mechanisms remains unclear given that kidney is one of the most vascularized organs which also express EnNaC and EC-SGK1. Further studies thus require closer examination on renal functions using the current experimental models as well as comparison among different vascular beds. Other limitations of the current study includes the inability to determine the temporal and causal relationship between blood pressure and arterial stiffening, which might require additional experimental groups with BP normalization.

### Perspective

Despite these limitations, this study uncovered a role for SGK1 in salt-sensitivity related vascular dysfunction (as assessed by changes in cellular stiffness) by utilizing a combination of mice with either global deletion or EC-selective deletion of SGK1 and complementary studies in primary cultured human aortic ECs. This study also presents a further downstream mechanism that may underlie SGK1 regulation of EC stiffening by showing aldosterone and salt-induced increases in actin polymerization were reduced by SGK1 inhibition in human aortic ECs. Yet, important questions remain regarding the mechanistic regulation of EnNaC by SGK1 in ECs and the role of EC-SGK1 in regulation of eNOS/NO signaling. Nevertheless, this study provides potential insights into approaches for developing therapeutic interventions for salt-sensitive hypertension-related cardiovascular dysfunction.

## NOVELTY AND RELEVANCE

### What Is New?

Upon DOCA-salt administration, mice with global serum and glucocorticoid regulated kinase 1 (SGK1) deletion had significantly reduced blood pressure, aortic endothelium stiffness and endothelial sodium channel activity as compared to littermate controls.

Selective endothelial cell (EC)-SGK1 deletion attenuated DOCA-salt-induced increases in blood pressure, EC and aortic stiffness.

SGK1 inhibitor, EMD638683, prevented aldosterone and high salt induced stiffening and actin polymerization in human aortic ECs.

### What Is Relevant?

It’s well documented that vascular stiffening contributes to overall morbidity and mortality rates of CVD. This study helps advance our understanding on vascular stiffening by uncovering the vital role of SGK1 in salt-sensitivity related vascular dysfunction and identifying a mechanistic link (through actin polymerization) between EC-SGK1 and vascular stiffening,

### Clinical/Pathophysiological Implications?

Endothelial dysfunction and vascular stiffening serve as early sign of various CVD, creating a window of opportunities for diagnostic and preventive intervention. Identification of cellular mechanisms underlying pathological cardiovascular stiffening hopefully provides the rationale for the targeting of therapeutic approaches for treating aspects of the CVD which impacts quality of life and mortality in a substantial portion of the population.

